# rSeqTU – a machine-learning based R package for prediction of bacterial transcription units

**DOI:** 10.1101/553057

**Authors:** Sheng-Yong Niu, Binqiang Liu, Qin Ma, Wen-Chi Chou

**Affiliations:** Department of Computer Science and Engineering, University of California San Diego, CA, USA; College of Mathematics and Physics, Qingdao University of Science and Technology, Qingdao, China; Biomedical Informatics, College of Medicine, The Ohio State University, Columbus, OH, USA; Infectious Disease and Microbiome Program, Broad Institute of MIT and Harvard, Cambridge, MA, USA

**Keywords:** machine-learning, bacteria, transcription unit, transcriptome

## Abstract

A transcription unit (TU) is composed of one or multiple adjacent genes on the same strand that are co-transcribed in mostly prokaryotes. Accurate identification of TUs is a crucial first step to delineate the transcriptional regulatory networks and elucidate the dynamic regulatory mechanisms encoded in various prokaryotic genomes. Many genomic features, e.g., gene intergenic distance, and transcriptomic features including continuous and stable RNA-seq reads count signals, have been collected from a large amount of experimental data and integrated into classification techniques to computationally predict genome-wide TUs. Although some tools and web servers are able to predict TUs based on bacterial RNA-seq data and genome sequences, there is a need to have an improved machine-learning prediction approach and a better comprehensive pipeline handling QC, TU prediction, and TU visualization. To enable users to efficiently perform TU identification on their local computers or high-performance clusters and provide a more accurate prediction, we develop an R package, named rSeqTU. rSeqTU uses a random forest algorithm to select essential features describing TUs and then uses support vector machine (SVM) to build TU prediction models. rSeqTU (available at https://s18692001.github.io/rSeqTU/) has six computational functionalities including read quality control, read mapping, training set generation, random-forest-based feature selection, TU prediction, and TU visualization.

## Introduction

The gene expression and regulation in bacteria use different machinery from eukaryotic organisms. Operon has been defined as a set of genes controlled by a single promoter are first co-transcribed into one mRNA molecule, and then the mRNA molecule is translated into multiple proteins (Jacob et al., 1960). Operationally, an operon uses a single promoter to regulate the set of genes. Functionally, the set of genes in the operon encodes proteins with related biological functions. The *lac* operon in *E. coli* is a typical operon that consists of a promoter, an operator, and three structural genes. The three genes, *lacZ, lacY*, and *lacA*, are co-transcribed into one mRNA transcript and subsequently translate into three proteins, β-galactosidase, β-galactoside permease, and Galactoside acetyltransferase. The *lac* operon is responsible for the transport and metabolism of lactose in many enteric bacteria. The discovery of the *lac* operon won the Nobel Prize in Physiology by Jacob and Monod in 1965 (Jacob et al., 1960).

Recently, many works revealed bacterial genes are not transcribed only in single operons but may be dynamically co-transcribed into mRNAs with different gene sets under different growth environments or conditions (Yan et al., 2018). Each of the co-transcribed gene set is called transcription units (TUs). The concept of TU is analogical to alternative spliced protein isoforms in eukaryotic systems that use different exons to produce protein isoforms. Although alternative splicing can use nonadjacent exons, a TU consists of a set of adjacent genes.

Several operon databases, such as RegulonDB (Santos-Zavaleta et al., 2019), MicrobesOnline(Dehal et al., 2010), and ProOpDB (Taboada et al., 2012) provide various levels of operon information describing genes only expressed in single TU or operon. While DOOR2(Mao et al., 2014) and OperomeDB (Chetal and Janga, 2015) provide the more comprehensive TUs describing genes are co-transcribed into different gene sets. Some TU or operon databases provide experiment-verified results while most of them rely on TU or operon predictions. Studies including DOOR2(Mao et al., 2014), SeqTU (Chou et al., 2015), and Rockhopper (McClure et al., 2013) use genomic information and gene expression profile to predict operon or TU with machine-learning and other approaches. Taboada et al. (Taboada et al., 2018) recently developed a new operon prediction method based on artificial neural network (ANN).

Other than in-silico prediction works, Yan et al., recently used SMRT-Cappable-seq and PacBio sequencing to re-examine the transcription units of *E. coli* grown under different conditions to provide a higher-resolution map of dynamic TUs (Yan et al., 2018). Yan et al.’s work revealed that TUs are better to describe the real bacterial transcription profiles and a gene can be co-transcribed into many different adjacent gene sets, TUs, under the same or different growth conditions. In our previous works (Chou et al., 2015) (Chen et al., 2017) we assumed a gene can only be co-transcribed into only one adjacent gene set, one TU. We also assumed cotranscribed gene pairs follow transitive relation, and we connect co-transcribed gene pairs into a larger gene sets to form a TU.

In this study, we focused on improving our machine-learning model for the prediction of the co-transcribed gene pairs and providing a user-friendly R package, rSeqTU, for a comprehensive pipeline including RNA-seq read analysis, TU prediction, and TU visualization.

## Results

In this rSeqTU R package, we updated the TU prediction model with random-forest-based feature selection and support vector machine (SVM). Besides, rSeqTU has a completed workflow performing RNA-seq read quality control (QC), RNA-seq read mapping, generation of TU results in two formats, and generation of IGV files for visualization (Figure 1).

**Figure 1.**
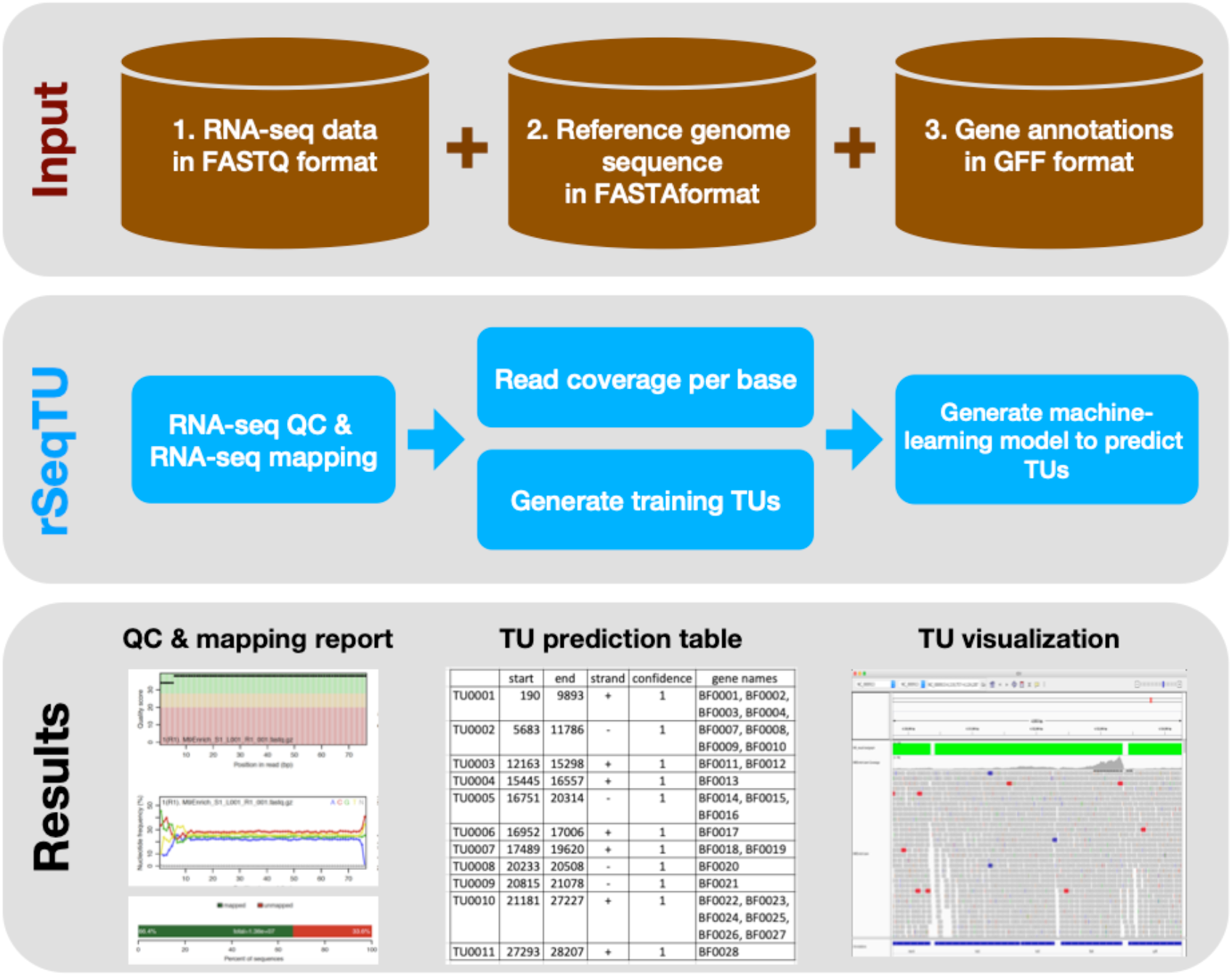
rSeqTU workflow uses input data to predict bacterial TUs. The rSeqTU workflow has three layers of schemas including input data, core processes, and major results. In the input data layer, rSeqTU needs RNA-seq data, reference genome sequence, and gene annotations. In the core process layer, rSeqTU performs QC, builds prediction models, and predict TUs. The results layer includes the QC and mapping results, TU prediction tables, and files for visualization in IGV.

rSeqTU requires three input data including RNA-seq data in FATSTQ format, reference genome sequence in FASTA format, and gene annotations in GFF format. With the input data, rSeqTU first performs RNA-seq data QC and RNA-seq read mapping to generate QC reports and mapping results in BAM format.

Then, rSeqTU uses whole-genome per-base read coverage and gene annotations to generate TU training data. The TU training data are co-transcribed gene pairs described by two sets of features generated based on an algorithm named SeqTU first presented by Chou et al. (Chou et al., 2015). rSeqTU then uses the TU training data to perform feature selection and build a TU prediction model. rSeqTU reports the prediction accuracy and uses the TU prediction model to identify all co-transcribed gene pairs in the given genome. rSeqTU outputs TU prediction results in single gene pairs and concatenated gene pairs. Last, rSeqTU converts TU results into IGV-compatible files for TU visualizations.

In short, rSeqTU produces RNA-seq read QC reports, RNA-seq mapping statistics and results, TU prediction results, and files for IGV visualization.

To evaluate rSeqTU R package, we used two sets of bacterial RNA-seq data of *Bacteroides fragilis (B. fragilis)* produced and published by Donaldson et al. [Donaldson et al. Science 2018]. These *B. fragilis* RNA-seq data were used to discover that human gut microbiome can use immunoglobulin A (IgA) to trigger robust host-microbial symbiosis for mucosal colonization. The study focused on investigating commensal colonization factors (CCFs), an operon, which were previously found to be essential for *B. fragilis* for colonization of colonic crypts [Lee et al. Nature 2013]. The CCF operon has five genes, ccfA-E, which are homologous to polysaccharide utilization systems, and the ccfA is activated by extracellular glycan sensing and is hypothesized to activate genes involved in mucosal colonization [Martens et al. JBC 2009]. To understand the function of *ccfA* gene, Donaldson et al. compared gene expression profiles beween *ccfA* overexpressed *B. fragilis* and wild-type *B. fragilis* and during laboratory culture growth. The RNA-seq data helped identify 24 out of 25 non-CCF genes that were differentially expressed and mapped to the biosynthesis loci for capsular polysaccharides A and C (PSA and PSC).

With the two RNA-seq data sets, reference genome sequence, and gene annotations, we performed a full run of rSeqTU analysis. The RNA-seq data QC and RNA-seq mapping were generated and shown in Figure 2.

**Figure 2.**
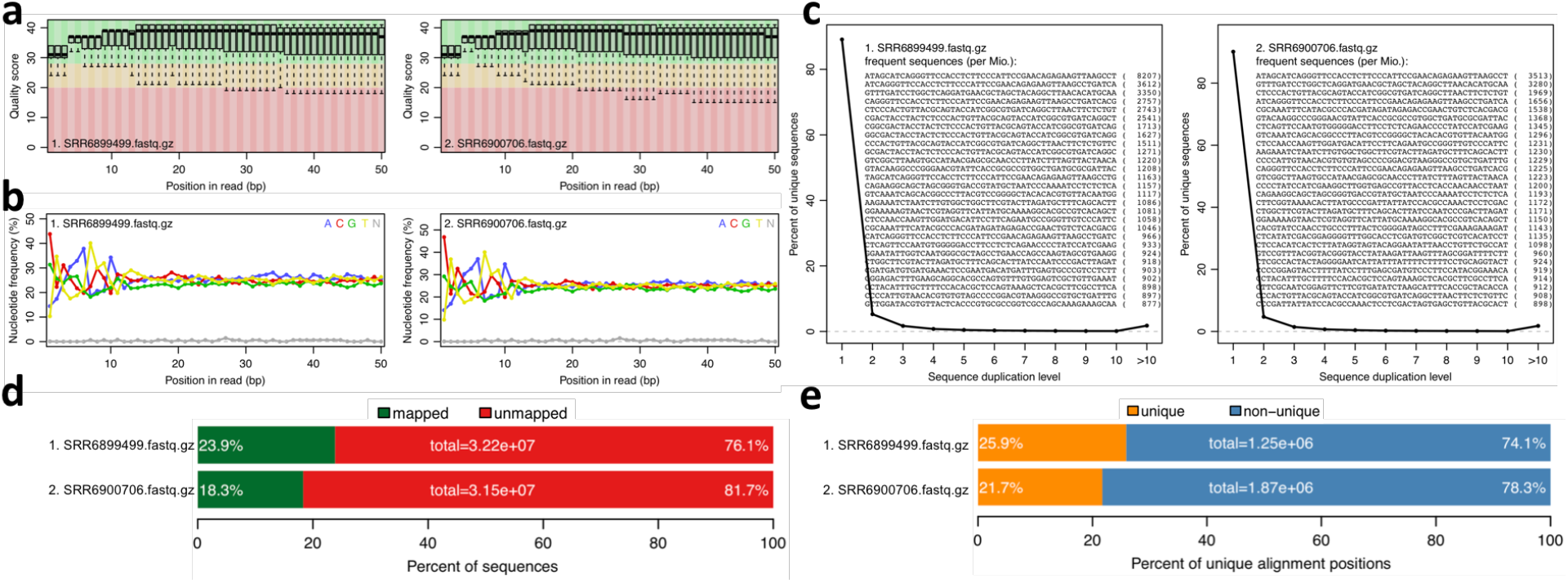
rSeqTU generates RNA-seq read QC reports and RNA-seq mapping statistics. SRR6899499 is *ccfA* overexpression data, and SRR6900706 is wild-type data. The panels a to e present quality scores, nucleotide frequency, sequence duplication, percentage of aligned bases plot, and percentage of unique and mapped reads.

In Figure 2, we generate QC report for both *ccfA* overexpression and wild-type. It shows the read quality score plot, which is good in general over 30 (Figure 2a). Also, it generated nucleotide frequency plot (Figure 2b), sequence duplication plot (Figure 2c), percentage of aligned bases plot (Figure 2d), and percentage of unique and mapped reads (Figure 2e). We could observe that the sequence duplication is not severe. The nucleotide frequency, aligned bases and mismatched bases information are in the normal range. The percentage of mapped reads and unique reads are lower than 30% as expected due to the most of the RNAs in the samples belong to mouse, the host, but not bacteria.

The two RNA-seq read mapping results were used to generate training data for TU prediction models, respectively. For *ccfA* overexpression data set, rSeqTU reported the sensitivity, specificity, and accuracy at 0.857, 0.999, and 0.963. For wild-type data set, rSeqTU reported the sensitivity, specificity, and accuracy at 0.885, 0.996, and 0.964. In general, we could find that rSeqTU generated high accuracy models after proper feature selections and cross-validation.

The two TU prediction models were used to predict co-transcribed gene pairs. There are 1759 and 1626 co-transcribed gene pairs predicted in *ccfA* overexpression and wild-type RNA-seq data sets. If we concatenated co-transcribed gene pairs, rSeqTU identified 2727 TUs including 2079 single-gene TUs, 271 two-gene TUs, and 377 TUs with more than two genes in *ccfA* overexpression RAN-seq data set. In wild-type RNA-seq data set, rSeqTU identified 2860 TUs including 2249 single-gene TUs, 256 two-gene TUs, and 355 TUs with more than two genes. rSeqTU then use the TU results to generate bedgraph files for the visualization in IGV (Figure 3). In the Figure 3, we showed a region of *B. fragilis* genome containing 8 genes. rSeqTU identified four TUs in the *ccfA* overexpression data (SRR6899499) and four TUs in the wild-type data (SRR6900706). However, the structure of the TUs are very different between two RNA-seq data sets. The two genes with locus tags, BF9343_RS17275 and BF9343_RS17280 were identified as a co-transcribed gene pair in the *ccfA* overexpression data but not in the wild-type data. The four genes with the locus tags, BF9343_RS17295, BF9343_RS17305, BF9343_RS17310, and BF9343_RS17315, were predicted as a single TU in the wild-type data but two TUs in the *ccfA* overexpression data.

**Figure 3.**
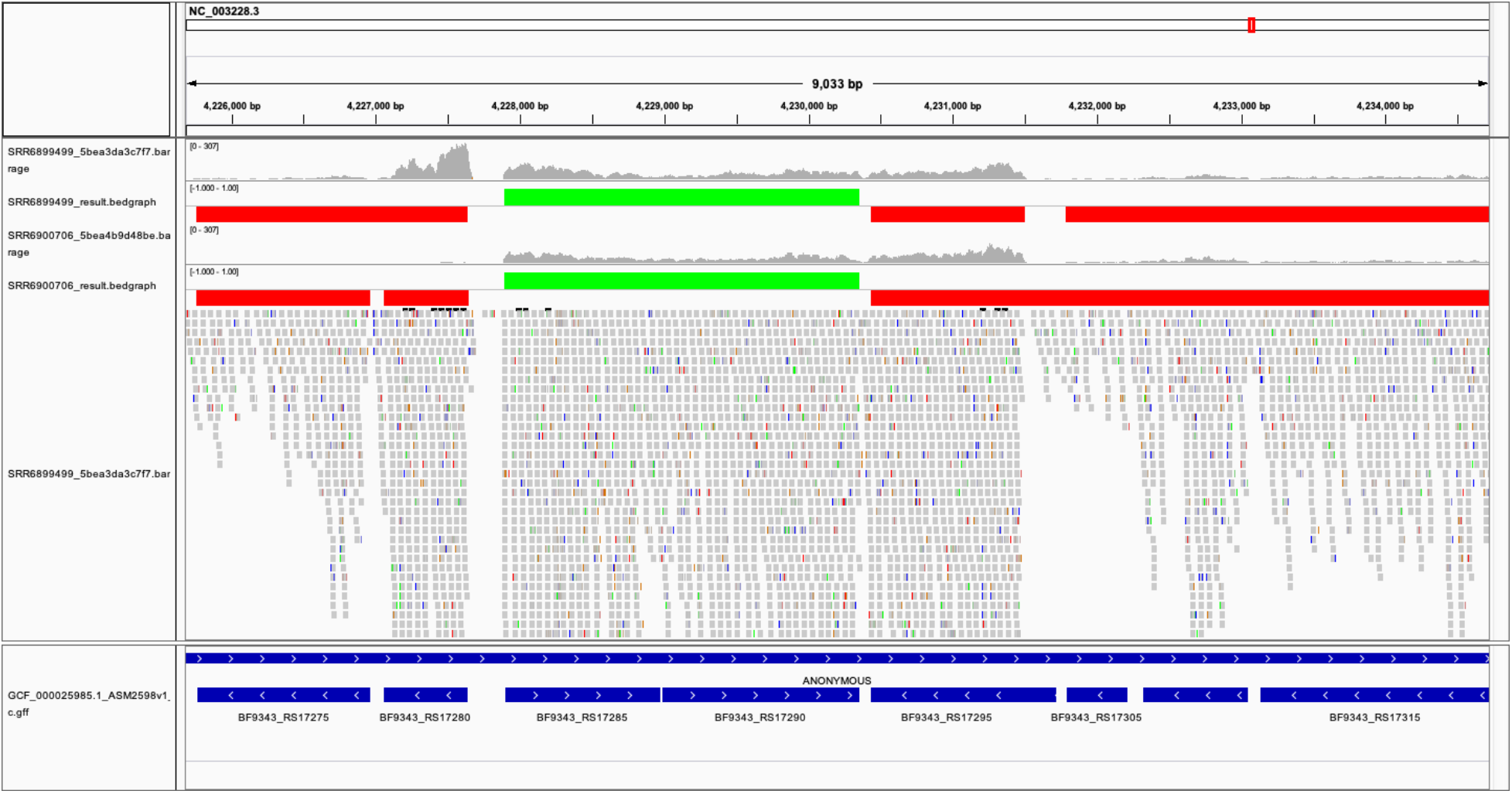
The visualization of predicted TUs on IGV. The predicted TUs are displayed in green and red bars for TUs on the forward and the reverse strands. The visualization also includes read coverage, gene annotations, and mapping results. SRR6899499 is *ccfA* overexpression data, and SRR6900706 is wild-type data.

## Material and Method

### New functions integrated or invented by rSeqTU

rSeqTU uses QuasR R pacakge to perform RNA-seq data QC and RNA-seq read mapping. The read mapping results are then processed by an algorithm named SeqTU first presented by Chou et al. (Chou et al., 2015). In brief, the SeqTU algorithm splits relatively long single genes into three parts including two sub-gene regions and an intergenic region, and then SeqTU uses RNA-seq per-base read coverage over the three parts to generate TU features to describe the continuity and stability of RNA-seq read coverage. SeqTU assumes the RNA-seq read coverage within a TU is continuous and stable like it is within a gene.

rSeqTU selects essential TU features by random forest-based and build TU prediction model by SVM using an R packages, Caret and e1071. rSeqTU converts TU prediction results into IGV-compatible files in bedgraph format for TU visualizations.

### *B. fragilis* RNA-seq data

We used two RNA-seq data from each triplicate experiment from NCBI’s SRA database with project accession number PRJNA445716. The accession numbers of the two data sets are SRR6899499 *(ccfA* overexpression) and SRR6900706 (wild-type). The reference genome sequence and gene annotations of Bacteroides fragilis NCTC 9343 are GCF_000025985.1_ASM2598v1_genomic.fna and GCF_000025985.1_ASM2598v1_genomic.gff.

## Discussion

rSeqTU is a machine learning-based R package for TU prediction, empowered by a random forest algorithm for feature selection and multiple graphical visualizations and interactive tables for customized downstream analysis. Its superior prediction performance has been demonstrated by testing multiple RNA-Seq datasets in *B. fragilis.* The source code and tutorial of rSeqTU is available at https://s18692001.github.io/rSeqTU/.

rSeqTU will be useful to understand transcriptional profiles of bacterial genomes in the gene level and the TU level. In addition to the single bacterium, rSeqTU may also be applied onto the metatranscriptomic data, the RNA-seq data of microbiome. The TUs of multiple bacteria may provide systemic view to understand how microbiome regulate functional translation and can be integrated with other metagenomic and metabolomic data (Niu et al., 2017).

A TU is dynamically composed by different adjacent genes under various conditions, and different TUs may overlap with each other under the same and different conditions. The dynamic TUs sharing the same gene(s) are called alternative transcription units (ATUs), and the identification of ATUs is recognized as a more challenging computational problem due to their condition-dependent nature. Meanwhile, the third-generation sequencing technology will shortly generate substantial genome-scale ATU datasets in the public domain for various prokaryotic organisms. Hence, advanced computational models are urgently needed for ATU prediction based on RNA-Seq data.

Intuitively, the output of rSeqTU can lay a solid foundation of ATU prediction as (i) a TU identified in our method can represents a maximal ATU clusters with apparent promoter and terminator and *(ii)* the TU can be used as an independent genomic region for further ATU prediction based on other genomic and transcriptomic features. If available, the ATUs along with related cis-regulatory motifs analysis will generate the dynamic regulatory networks in a bacterial genome to a higher resolution and an advanced level.

## Funding

This work was supported by Dr. Qin Ma’s startup funding in the Department of Biomedical Informatics at the Ohio State University. This work used the Extreme Science and Engineering Discovery Environment (XSEDE), which is supported by the National Science Foundation #ACI-1548562. The content is solely the responsibility of the authors and does not necessarily represent the official views of the National Science Foundation.

## Conflict of interest statement

None declared.

